## Supplementary material for "Is adaptation limited by mutation? A timescale-dependent effect of genetic diversity on the adaptive substitution rate in animals": All supplementary material

| Species | Bioproject | data_type | #individuals | publication | Reference genome |
|---|---|---|---|---|---|
| *Gorilla gorilla* | PRJNA189439 | Genome | 20 | Prado-Martinez et al. 2013 | Ensembl (release 89) |
| *Homo sapiens* | PRJEB8350 | Exome | 19 | Teixeira et al. 2015 | Ensembl (release 89) |
| *Pan troglodytes* | PRJEB8350 | Exome | 20 | Teixeira et al. 2015 | Ensembl (release 89) |
| *Papio anubis* | PRJNA54005 | Genome | 5 | unpublished baboon genome project | Ensembl (release 89) |
| *Pongo abelii* | PRJNA189439 and PRJEB1675 | Genome | 10 | Prado-Martinez et al. 2013 | Ensembl (release 89) |
| *Macaca mulatta* | PRJNA251548 | Exome | 20 | Xue et al. 2016 | Ensembl (release 89) |
| *Meleagris gallopavo* | PRJNA271731 | RNA-seq | 10 | Wright et al. 2015 | NA |
| *Phasianus colchicus* | PRJNA271731 | RNA-seq | 11 | Wright et al. 2015 | NA |
| *Pavo cristatus* | PRJNA271731 | RNA-seq | 10 | Wright et al. 2015 | NA |
| *Numida meleagris* | PRJNA271731 | RNA-seq | 7 | Wright et al. 2015 | NA |
| *Anas platyrhynchos* | PRJNA271731 | RNA-seq | 10 | Wright et al. 2015 | NA |
| *Anser cygnoides* | PRJNA271731 | RNA-seq | 10 | Wright et al. 2015 | NA |
| *Ficedula albicollis* | PRJEB2984 | Genome | 20 | Ellegren et al. 2012 | NCBI FicAlb1.5 |
| *Geospiza difficilis* | PRJNA263122 | Genome | 8 | Lamichhaney et al. 2015 | NCBI Geofor1.0 |
| *Parus major* | PRJNA381923 | Genome | 10 | Corcoran et al. 2017 | //ftp.ncbi.nlm.nih.gov/genomes/all/GCF/001/522/545/GCF_001522545.2_Parus_major1.1/GCF_001522545.2_Parus_major1.1_genomic.gff.gz |
| *Corvus sp.* | PRJEB9057 | Genome | 10 | Vijay et al. 2017 | //ftp.ncbi.nlm.nih.gov/genomes/all/GCF/000/738/735/GCF_000738735.2_ASM73873v2/GCF_000738735.2_ASM73873v2_genomic.gff.gz |
| *Taniopygia guttata* | PRJEB10586 | Genome | 20 | Singhal et al. 2016 | //ftp.ncbi.nlm.nih.gov/genomes/all/GCF/000/151/805/GCF_000151805.1_Taeniopygia_guttata-3.2.4/GCF_000151805.1_Taeniopygia_guttata-3.2.4_genomic.gff.gz |
| *Maniola jurtina* | PRJNA530965 | target capture | 20 | newly generated | NA |
| *Melanargia galathea* | PRJNA530965 | target capture | 10 | newly generated | NA |
| *Aphantopus hyperantus* | PRJNA530965 | target capture | 7 | newly generated | NA |
| *Pyronia tithonus* | PRJNA530965 | target capture | 7 | newly generated | NA |
| *Pyronia bathseba* | PRJNA530965 | target capture | 8 | newly generated | NA |

| | | | | | |
|---|---|---|---|---|---|
| *Formica sanguinea* | PRJNA530965 | target capture | 10 | newly generated | NA |
| *Formica cunicularia* | PRJNA530965 | target capture | 6 | newly generated | NA |
| *Formica pratensis* | PRJNA530965 | target capture | 8 | newly generated | NA |
| *Formica fusca* | PRJNA530965 | target capture | 8 | newly generated | NA |
| *Allolobophora chlorotica L1* | PRJNA530965 | target capture | 19 | newly generated | NA |
| *Allolobophora chlorotica L2* | PRJNA530965 | target capture | 8 | newly generated | NA |
| *Allolobophora chlorotica L4* | PRJNA530965 | target capture | 9 | newly generated | NA |
| *Aporrectodea icterica* | PRJNA530965 | target capture | 10 | newly generated | NA |
| *lumbricus terrestris* | PRJNA530965 | target capture | 9 | newly generated | NA |
| *Lineus lacteus* | PRJNA530965 | target capture | 9 | newly generated | NA |
| *Lineus sanguineus* | PRJNA530965 | target capture | 9 | newly generated | NA |
| *Lineus longissimus* | PRJNA530965 | target capture | 6 | newly generated | NA |
| *Lineus ruber* | PRJNA530965 | target capture | 8 | newly generated | NA |
| *Mytilus galloprovincialis* | PRJNA530965 | target capture | 9 | newly generated | NA |
| *Mytilus californianus* | PRJNA530965 | target capture | 16 | newly generated | NA |
| *Mytilus edulis* | PRJNA530965 | target capture | 10 | newly generated | NA |
| *Mytilus trossulus* | PRJNA530965 | target capture | 10 | newly generated | NA |
| *Drosophila melanogaster* | SRP006733 | Genome | 10 | Pool et al. 2012 | //ftp.ncbi.nlm.nih.gov/genomes/all/GCF/000/001/215/GCF_000001215.4_Release_6_plus_ISO1_MT/GCF_000001215.4_Release_6_plus_ISO1_MT_genomic.fna.gz |
| *Drosophila sechellia* | PRJNA395473 | Genome | 8 | Schrider et al. 2018 | ftp://ftp.ncbi.nlm.nih.gov/genomes/all/GCF/000/005/215/GCF_000005215.3_dsec_caf1/GCF_000005215.3_dsec_caf1_genomic.fna.gz |
| *Drosophila simulans* | PRJNA215932 | Genome | 10 | Rogers et al. 2014 | ftp://ftp.ncbi.nlm.nih.gov/genomes/all/GCF/000/754/195/GCF_000754195.2_ASM75419v2/GCF_000754195.2_ASM75419v2_genomic.fna.gz |
| *Drosophila santomea* | PRJNA395473 | Genome | 17 | Turissini & Matute 2017 | ftp://ftp.ncbi.nlm.nih.gov/genomes/all/GCF/000/005/975/GCF_000005975.2_dyak_caf1/GCF_000005975.2_dyak_caf1_genomic.fna.gz |
| *Drosophila yakuba* | PRJNA395473 | Genome | 20 | Turissini & Matute 2017 | ftp://ftp.ncbi.nlm.nih.gov/genomes/all/GCF/000/005/975/GCF_000005975.2_dyak_caf1/GCF_000005975.2_dyak_caf1_genomic.fna.gz |
| *Drosophila teissieri* | PRJNA395473 | Genome | 11 | Turissini & Matute 2017 | ftp://ftp.ncbi.nlm.nih.gov/genomes/all/GCF/000/005/975/GCF_000005975.2_dyak_caf1/GCF_000005975.2_dyak_caf1_genomic.fna.gz |

| | | | | | |
|---|---|---|---|---|---|
| *Mus musculus castaneus* | PRJEB2176 | Genome | 10 | Harr et al. 2016 | ftp://ftp.ncbi.nlm.nih.gov/genomes/all/GCF/000/001/635/GCF_000001635.26_GRCm38.p6/GCF_000001635.26_GRCm38.p6_genomic.fna.gz |
| *Mus spretus* | PRJEB11742 | Genome | 8 | Harr et al. 2016 | ftp://ftp.ncbi.nlm.nih.gov/genomes/all/GCA/001/624/865/GCA_001624865.1_SPRET_EiJ_v1/GCA_001624865.1_SPRET_EiJ_v1_genomic.fna.gz |
| *Rattus norvegicus* | PRJEB2922 | Genome | 12 | Deinum et al. 2015 | ftp://ftp.ncbi.nlm.nih.gov/genomes/all/GCF/000/001/895/GCF_000001895.5_Rnor_6.0/GCF_000001895.5_Rnor_6.0_genomic.fna.gz |
| *Microtus ochrogaster* | PRJNA428754 | RNA-seq | 18 | NA | NA |
| *Microtus arvalis* | PRJNA249058 | RNA-seq | 7 | Romiguier et al. 2014 | NA |

**Table S1 : Details of the species used in this study and numbers of individuals for each species.**

| Taxonomic group | minimum orthogroup number | maximum orthogroup number |
|---|---|---|
| ants | 1249 | 1261 |
| butterflies | 1561 | 1671 |
| earth worms | 796 | 1460 |
| Ribbon worms | 1049 | 1413 |
| mussles | 1525 | 1525 |
| primates | 8604 | 8604 |
| passerines | 6755 | 6755 |
| fowls | 4439 | 4439 |
| rodents | 3994 | 3994 |
| flies | 7900 | 7900 |

**Table S2: Number of orthogroups for each taxonomic group.**

The differences in terms of number of orthogroups comes from the fact that we not only kept orthogroups with all species but also orthogroups with all species but one to estimate dN/dS value for each terminal branches in order to maximize the number of substitutions for data sets generated by exon capture.

| species | Number of indivduals | Chosen sample size | # non-synonymous SNPs | # synonymous SNPs | # SNPs total | #GC-cons, Non-synonymous SNPs | #GC-cons, Synonymous SNPs | #GC-cons, SNPs total |
|---|---|---|---|---|---|---|---|---|
| *F. fusca* | 8 | 10 | 4278 | 6578 | 10856 | 270 | 166 | 436 |
| *F. sanguinea* | 10 | 16 | 3242 | 5343 | 8585 | 363 | 224 | 587 |
| *F. cunicularia* | 6 | 8 | 4035 | 6355 | 10390 | 212 | 121 | 333 |
| *F. pratensis* | 8 | 12 | 1773 | 2235 | 4008 | 191 | 87 | 278 |
| *M. galathea* | 10 | 10 | 1190 | 3309 | 4499 | 621 | 346 | 966 |
| *M. jurtina* | 20 | 32 | 7310 | 15926 | 23235 | 1863 | 2036 | 3898 |
| *A. hyperanthus* | 7 | 8 | 1441 | 2385 | 3826 | 734 | 350 | 1084 |
| *P. tithonus* | 7 | 10 | 1534 | 2372 | 3906 | 369 | 278 | 647 |
| *P. bathseba* | 8 | 10 | 1611 | 2676 | 4287 | 395 | 315 | 710 |
| *M. californianus* | 16 | 24 | 6370 | 13436 | 19806 | 1467 | 2209 | 3675 |
| *M. trossulus* | 10 | 14 | 6329 | 18994 | 25323 | 1665 | 3382 | 5047 |
| *M. galloprovincialis* | 9 | 12 | 3298 | 9089 | 12387 | 839 | 1665 | 2504 |
| *M. edulis* | 10 | 12 | 5497 | 15987 | 21485 | 1326 | 2590 | 3915 |
| *A. chlorotica L1* | 19 | 26 | 1751 | 3094 | 4845 | 472 | 442 | 914 |
| *A. chlorotica L2* | 8 | 8 | 350 | 554 | 904 | 69 | 56 | 126 |
| *A. chlorotica L4* | 9 | 12 | 3562 | 7895 | 11457 | 701 | 808 | 1508 |
| *A. icterica* | 10 | 12 | 1657 | 3778 | 5435 | 360 | 387 | 747 |
| *L. terrestris* | 9 | 8 | 238 | 940 | 1178 | 38 | 77 | 115 |
| *L. lacteus* | 9 | 12 | 5896 | 20421 | 26317 | 1292 | 2495 | 3788 |
| *L. longissimus* | 6 | 8 | 55 | 99 | 154 | 11 | 7 | 18 |
| *L. sanguineus* | 9 | 10 | 954 | 3248 | 4202 | 192 | 397 | 589 |
| *L. ruber* | 8 | 12 | 1028 | 1691 | 2718 | 196 | 168 | 364 |
| *H. sapiens* | 19 | 28 | 2281 | 2812 | 5093 | 264 | 123 | 387 |
| *P. troglodytes* | 20 | 30 | 4744 | 6558 | 11302 | 559 | 266 | 825 |
| *G. gorilla* | 20 | 30 | 3649 | 4842 | 8491 | 370 | 180 | 550 |
| *P. anubis* | 5 | 8 | 2038 | 4006 | 6044 | 242 | 164 | 407 |
| *P. abelii* | 10 | 16 | 5687 | 9300 | 14987 | 594 | 360 | 954 |
| *M. mulatta* | 19 | 28 | 5257 | 9976 | 15232 | 608 | 560 | 1169 |
| *A. pltyrhynchos* | 10 | 16 | 3796 | 13089 | 16884 | 206 | 438 | 645 |
| *A. cygnoides* | 10 | 14 | 1296 | 4824 | 6119 | 103 | 127 | 230 |
| *M. gallopavo* | 10 | 10 | 1005 | 3791 | 4795 | 61 | 52 | 113 |
| *N. meleagris* | 10 | 16 | 1319 | 6241 | 7560 | 90 | 135 | 224 |
| *P. cristatus* | 10 | 14 | 602 | 2214 | 2816 | 48 | 51 | 99 |
| *P. colchicus* | 11 | 14 | 3051 | 9703 | 12754 | 172 | 172 | 343 |
| *P. major* | 10 | 16 | 6906 | 14414 | 21320 | 2702 | 2292 | 4994 |
| *F. albicolis* | 20 | 16 | 16242 | 20681 | 36923 | 2818 | 2224 | 5042 |
| *Corvus sp.* | 10 | 14 | 817 | 1355 | 2172 | 115 | 83 | 197 |
| *G. difficilis* | 8 | 10 | 2090 | 3280 | 5370 | 374 | 232 | 606 |
| *T. guttata* | 20 | 16 | 36318 | 102249 | 138567 | 6588 | 7775 | 14363 |
| *R. norvegicus* | 12 | 18 | 5023 | 10224 | 15246 | 645 | 574 | 1220 |
| *M. arvalis* | 7 | 10 | 1783 | 6351 | 8134 | 232 | 377 | 609 |
| *M. ochrogaster* | 18 | 18 | 2455 | 5340 | 7795 | 448 | 354 | 801 |
| *M. spretus* | 8 | 12 | 58687 | 14289 | 72976 | 812 | 921 | 1733 |
| *M. m. castaneus* | 10 | 12 | 5524 | 17996 | 23520 | 778 | 1068 | 1846 |
| *D. melanogaster* | 10 | 16 | 36213 | 138306 | 174518 | 9574 | 15862 | 25436 |
| *D. teissieri* | 11 | 18 | 48565 | 247605 | 296171 | 14259 | 35930 | 50188 |
| *D. santomea* | 17 | 28 | 16104 | 54878 | 70982 | 2374 | 4142 | 6516 |
| *D. yakuba* | 20 | 12 | 37921 | 164003 | 201924 | 11090 | 22635 | 33725 |
| *D. simulans* | 10 | 16 | 74864 | 352964 | 427828 | 21397 | 47094 | 68490 |
| *D. sechellia* | 8 | 12 | 379 | 591 | 970 | 92 | 61 | 153 |

**Table S3: SNPs counts for each species.**

| taxonomic group | tree topologies references |
|---|---|
| Catarrhine primates | Perelman P, Johnson WE, Roos C, Seuánez HN, Horvath JE, Moreira MA, Kessing B, Pontius J, Roelke M, Rumpler Y, Schneider MP. 2011. A molecular phylogeny of living primates. PLoS genetics. 7(3):e1001342. |
| Galloanserae | Wright AE, Harrison PW, Zimmer F, Montgomery SH, Pointer MA, Mank JE. 2015. Variation in promiscuity and sexual selection drives avian rate of Faster-Z evolution. Molecular ecology. 24(6):1218-35. |
| Passeriformes | Barker FK, Barrowclough GF, Groth JG. 2002. A phylogenetic hypothesis for passerine birds: taxonomic and biogeographic implications of an analysis of nuclear DNA sequence data. Proceedings of the Royal Society B: Biological Sciences 269:295–308. |
| Muroidea | Steppan SJ, Adkins RM, Anderson J, Thorne J. 2004. Phylogeny and Divergence-Date Estimates of Rapid Radiations in Muroid Rodents Based on Multiple Nuclear Genes. Systematic Biology 53:533–553. |
| Mussels | Distel DL. 2000. Phylogenetic Relationships among Mytilidae (Bivalvia): 18S rRNA Data Suggest Convergence in Mytilid Body Plans. Molecular Phylogenetics and Evolution 15:25–33. |
| Satyrinae butterflies | Peña C, Wahlberg N, Weingartner E, Kodandaramaiah U, Nylin S, Freitas AV, Brower AV. 2006. Higher level phylogeny of Satyrinae butterflies (Lepidoptera: Nymphalidae) based on DNA sequence data. Molecular phylogenetics and evolution. 40(1):29-49. |
| Formica ants | J. Romiguier, J. Rolland, C. Morandin, L. Keller. 2018. Phylogenomics of palearctic Formica species suggests a single origin of temporary parasitism and gives insights to the evolutionary pathway toward slave-making behaviour. BMC evolutionary biology 18(1):40. |
| Earth worms | R. A. King, A. L. Tibble, W. O. C. Symondson. 2008. Opening a can of worms: unprecedented sympatric cryptic diversity within British lumbricid earthworms. *Molecular Ecology* 17, 4684-4698. |
| Nemertea | Thollesson, Mikael, and Jon L. Norenburg. 2003. Ribbon worm relationships: a phylogeny of the phylum Nemertea. Proceedings of the Royal Society of London B: Biological Sciences 27(1513): 407-415. |
| Drosophila | Obbard DJ, Maclennan J, Kim KW, Rambaut A, O'grady PM, Jiggins FM. 2012. Estimating divergence dates and substitution rates in the Drosophila phylogeny. Molecular Biology and Evolution. 29(11):3459-73. |

**Table S4 : Sources of the tree topologies of each taxonomic group used to estimate branch length and map substitutions.**

| Species | Propagule size (cm) | source |
|---|---|---|
| *Formica fusca* | 14 | Forel 1890 |
| *Formica sanguinea* | 10 | Forel 1909 |
| *Formica cunicularia* | 8.5 | Collingwood 1979 |
| *Formica pratensis* | 10.4 | Bolton 1995 |
| *Melanargia galathea* | 0.102 | García-Barros 2000 |
| *Maniola jurtina* | 0.0535 | García-Barros 2000 |
| *Aphantopus hyperantus* | 0.0792 | García-Barros 2000 |
| *Pyronia tithonus* | 0.0628 | García-Barros 2000 |
| *Pyronia bathseba* | 0.0802 | García-Barros 2000 |
| *Mytilus californianus* | 0.01 | Bayne et al. 1983 |
| *Mytilus trossulus* | 0.01 | Bayne et al. 1983 |
| *Mytilus galloprovincialis* | 0.01 | Bayne et al. 1983 |
| *Mytilus edulis* | 0.01 | Bayne et al. 1983 |
| *Allolobophora chlorotica L1* | 0.0238 | Eijsackers 2011 |
| *Allolobophora chlorotica L2* | 0.0238 | Eijsackers 2011 |
| *Allolobophora chlorotica L4* | 0.0238 | Eijsackers 2011 |
| *Aporrecta icterica* | 0.0411 | Eijsackers 2011 |
| *Lumbricus terrestris* | 0.5 | Cloudsley-Thompson and Sankey 1961 |
| *Lineus lacteus* | 0.02 | Bierne, 1983 |
| *Lineus longissimus* | 0.02 | Bierne, 1983 |
| *Lineus sanguineaus* | 0.02 | Bierne, 1983 |
| *Lineus ruber* | 0.5 | Bierne, 1983 |
| *Homo sapiens* | 93.4 | De Magalhaes and Costa 2009 |
| *Pan troglodytes* | 45.12 | De Magalhaes and Costa 2009 |
| *Gorilla gorilla* | 78 | De Magalhaes and Costa 2009 |
| *Papio anubis* | 55.95 | De Magalhaes and Costa 2009 |
| *Pongo abelii* | 42.48 | De Magalhaes and Costa 2009 |
| *Macaca mulatta* | 31.1 | De Magalhaes and Costa 2009 |
| *Anas platyrhynchos* | 55 | Del Hoyo et al. 1992 |
| *Anser cygnoides* | 87 | Del Hoyo et al. 1992 |
| *Meleagris gallopavo* | 90 | Del Hoyo et al. 1992 |
| *Numida meleagris* | 53 | Del Hoyo et al. 1992 |
| *Pavo cristatus* | 95 | Del Hoyo et al. 1992 |
| *Phasianus colchicus* | 57.5 | Del Hoyo et al. 1992 |

| | | |
|---|---|---|
| *Parus major* | 13.5 | Del Hoyo et al. 1992 |
| *Ficedula albicollis* | 13 | Del Hoyo et al. 1992 |
| *Corvus sp.* | 50.5 | Del Hoyo et al. 1992 |
| *Geospiza difficilis* | 11.5 | Del Hoyo et al. 1992 |
| *Taniopygia guttata* | 10 | Del Hoyo et al. 1992 |
| *Rattus norvegicus* | 13.717 | De Magalhaes and Costa 2009 |
| *Microtus arvalis* | 7.184 | De Magalhaes and Costa 2009 |
| *Microtus ochrogaster* | 10.6 | De Magalhaes and Costa 2009 |
| *Mus musculus musculus* | 6.06 | De Magalhaes and Costa 2009 |
| *Mus spretus* | 6.37 | Inferred from De Magalhaes and Costa 2009 |
| *Drosophila melanogaster* | 0.0525 | Lott et al. 2007 |
| *Drosophila teissieri* | NA | NA |
| *Drosophila santomea* | NA | NA |
| *Drosophila yakuba* | 0.0475 | Lott et al. 2007 |
| *Drosophila simulans* | 0.05 | Lott et al. 2007 |
| *Drosophila sechellia* | 0.06 | Lott et al. 2007 |

| **Species** | **adult size (cm)** | **source** |
|---|---|---|
| *Formica fusca* | 14 | Forel 1890 |
| *Formica sanguinea* | 10 | Forel 1909 |
| *Formica cunicularia* | 8.5 | Collingwood 1979 |
| *Formica pratensis* | 10.4 | Bolton 1995 |
| *Melanargia galathea* | 2.59 | García-Barros 2000 |
| *Maniola jurtina* | 2.58 | García-Barros 2000 |
| *Aphantopus hyperantus* | 2.12 | García-Barros 2000 |
| *Pyronia tithonus* | 1.89 | García-Barros 2000 |
| *Pyronia bathseba* | 2.04 | García-Barros 2000 |
| *Mytilus californianus* | 7.5 | MArine Life Information Network, 2006 |
| *Mytilus trossulus* | 7.5 | MArine Life Information Network, 2006 |
| *Mytilus galloprovincialis* | 7.5 | MArine Life Information Network, 2006 |
| *Mytilus edulis* | 7.5 | MArine Life Information Network, 2006 |
| *Allolobophora chlorotica L1* | 5.5 | The Trustees of the Natural |

| | | |
|---|---|---|
| | | History Museum,2010 |
| *Allolobophora chlorotica L2* | 5.5 | The Trustees of the Natural History Museum,2010 |
| *Allolobophora chlorotica L4* | 5.5 | The Trustees of the Natural History Museum,2010 |
| *Aporrecta icterica* | 9.5 | Sims and Gerard 1985 |
| *Lumbricus terrestris* | 25 | Cloudsley-Thompson and Sankey 1961 |
| *Lineus lacteus* | 17.5 | Gontcharoff 1951 |
| *Lineus longissimus* | 1000 | Gontcharoff 1951 |
| *Lineus sanguineus* | NA | NA |
| *Lineus ruber* | 5 | Gontcharoff 1951, Bierne 1970 |
| *Homo sapiens* | 163 | Ogden et al. 2004 |
| *Pan troglodytes* | 79.6 | Jones et al. 2009 |
| *Gorilla gorilla* | 137.5 | Wood 1979 |
| *Papio anubis* | 85 | Fleagle 2013 |
| *Pongo abelii* | 83 | Groves 1971 |
| *Macaca mulatta* | 55.5 | Jones et al. 2009 |
| *Anas platyrhynchos* | 55 | Del Hoyo et al. 1992 |
| *Anser cygnoides* | 87 | Del Hoyo et al. 1992 |
| *Meleagris gallopavo* | 90 | Del Hoyo et al. 1992 |
| *Numida meleagris* | 53 | Del Hoyo et al. 1992 |
| *Pavo cristatus* | 95 | Del Hoyo et al. 1992 |
| *Phasianus colchicus* | 57.5 | Del Hoyo et al. 1992 |
| *Parus major* | 13.5 | Del Hoyo et al. 1992 |
| *Ficedula albicollis* | 13 | Del Hoyo et al. 1992 |
| *Corvus sp.* | 50.5 | Del Hoyo et al. 1992 |
| *Geospiza difficilis* | 11.5 | Del Hoyo et al. 1992 |
| *Taniopygia guttata* | 10 | Del Hoyo et al. 1992 |
| *Rattus norvegicus* | 21.5 | Burton and Burton 2002 |
| *Microtus arvalis* | 11.1 | Jones et al. 2009 |
| *Microtus ochrogaster* | 15.2 | Jones et al. 2009 |
| *Mus musculus musculus* | 8 | Berry 1970 |
| *Mus spretus* | 8.6 | Palomo et al. 2009 |
| *Drosophila melanogaster* | 0.85 | Pitnick et al. 2002 |
| *Drosophila teissieri* | NA | NA |
| *Drosophila santomea* | NA | NA |

| | | |
|---|---|---|
| *Drosophila yakuba* | NA | NA |
| *Drosophila simulans* | 1.03 | NA |
| *Drosophila sechellia* | NA | NA |

| **Species** | **body mass (g)** | **source** |
|---|---|---|
| *Formica fusca* | NA | NA |
| *Formica sanguinea* | NA | NA |
| *Formica cunicularia* | NA | NA |
| *Formica pratensis* | 0.0119 | Keller & Passera, 1989 |
| *Melanargia galathea* | NA | NA |
| *Maniola jurtina* | 0.05 | Svärd &Wiklund,1989 |
| *Aphantopus hyperantus* | 0.0376 | Svärd &Wiklund,1989 |
| *Pyronia tithonus* | 0,04 | Corbet, 2000 |
| *Pyronia bathseba* | NA | NA |
| *Mytilus californianus* | 37.5 | MArine Life Information Network, 2006 |
| *Mytilus trossulus* | 37.5 | MArine Life Information Network 2006 |
| *Mytilus galloprovincialis* | 37.5 | MArine Life Information Network, 2006 |
| *Mytilus edulis* | 37.5 | MArine Life Information Network, 2006 |
| *Allolobophora chlorotica L1* | 0.3 | Butt 1997 |
| *Allolobophora chlorotica L2* | 0.3 | Butt 1997 |
| *Allolobophora chlorotica L4* | 0.3 | Butt 1997 |
| *Aporrecta icterica* | 0.95 | Bouché 1972 |
| *Lumbricus terrestris* | 7.5 | Quillin, 1999 |
| *Lineus lacteus* | NA | NA |
| *Lineus longissimus* | NA | NA |
| *Lineus sanguineaus* | NA | NA |
| *Lineus ruber* | NA | NA |
| *Homo sapiens* | 62000 | De Magalhaes and Costa 2009 |
| *Pan troglodytes* | 45000 | De Magalhaes and Costa 2009 |
| *Gorilla gorilla* | 93000 | De Magalhaes and Costa 2009 |
| *Papio anubis* | 14700 | De Magalhaes and Costa 2009 |
| *Pongo abelii* | 45000 | De Magalhaes and Costa 2009 |
| *Macaca mulatta* | 8240 | De Magalhaes and Costa 2009 |

| | | |
|---|---|---|
| *Anas platyrhynchos* | 1027 | De Magalhaes and Costa 2009 |
| *Anser cygnoides* | 3150 | De Magalhaes and Costa 2009 |
| *Meleagris gallopavo* | 4000 | De Magalhaes and Costa 2009 |
| *Numida meleagris* | 1479 | De Magalhaes and Costa 2009 |
| *Pavo cristatus* | 3375 | De Magalhaes and Costa 2009 |
| *Phasianus colchicus* | 999 | De Magalhaes and Costa 2009 |
| *Parus major* | 17 | Del Hoyo et al. 1992 |
| *Ficedula albicollis* | 12 | Del Hoyo et al. 1992 |
| *Corvus sp.* | 499 | Del Hoyo et al. 1992 |
| *Geospiza difficilis* | 16.15 | Del Hoyo et al. 1992 |
| *Taniopygia guttata* | 10 | Del Hoyo et al. 1992 |
| *Rattus norvegicus* | 320 | De Magalhaes and Costa 2009 |
| *Microtus arvalis* | 27.5 | De Magalhaes and Costa 2009 |
| *Microtus ochrogaster* | 50 | De Magalhaes and Costa 2009 |
| *Mus musculus musculus* | 20.5 | De Magalhaes and Costa 2009 |
| *Mus spretus* | 17 | Palomo et al. 2009 |
| *Drosophila melanogaster* | 0.00115 | Klok et al. 2009 |
| *Drosophila teissieri* | NA | NA |
| *Drosophila santomea* | NA | NA |
| *Drosophila yakuba* | NA | NA |
| *Drosophila simulans* | NA | NA |
| *Drosophila sechellia* | NA | NA |

| **Species** | **Fecundity (number of offspring per year)** | **source** |
|---|---|---|
| *Formica fusca* | NA | NA |
| *Formica sanguinea* | NA | NA |
| *Formica cunicularia* | NA | NA |
| *Formica pratensis* | NA | NA |
| *Melanargia galathea* | NA | NA |
| *Maniola jurtina* | NA | NA |
| *Aphantopus hyperantus* | 140 | Lafranchis et al. 2015 |
| *Pyronia tithonus* | 125 | Lafranchis et al. 2015 |
| *Pyronia bathseba* | NA | NA |
| *Mytilus californianus* | 110000 | MArine Life Information |

| | | |
|---|---|---|
| | | Network, 2006 |
| *Mytilus trossulus* | 110000 | MArine Life Information Network, 2006 |
| *Mytilus galloprovincialis* | 110000 | MArine Life Information Network, 2006 |
| *Mytilus edulis* | 110000 | MArine Life Information Network, 2006 |
| *Allolobophora chlorotica L1* | 0.74 | Edwards & Bohlen 1996 |
| *Allolobophora chlorotica L2* | 0.74 | Edwards & Bohlen 1996 |
| *Allolobophora chlorotica L4* | 0.74 | Edwards & Bohlen 1996 |
| *Aporrecta icterica* | 2.67 | Booth et al. 2000 |
| *Lumbricus terrestris* | NA | NA |
| *Lineus lacteus* | NA | NA |
| *Lineus longissimus* | NA | NA |
| *Lineus sanguineaus* | NA | NA |
| *Lineus ruber* | NA | NA |
| *Homo sapiens* | 0.0008219178 | De Magalhaes and Costa 2009 |
| *Pan troglodytes* | 0.0005479452 | De Magalhaes and Costa 2009 |
| *Gorilla gorilla* | 0.0008219178 | De Magalhaes and Costa 2009 |
| *Papio anubis* | 0.002191781 | De Magalhaes and Costa 2009 |
| *Pongo abelii* | 0.0005479452 | De Magalhaes and Costa 2009 |
| *Macaca mulatta* | 0.002739726 | De Magalhaes and Costa 2009 |
| *Anas platyrhynchos* | 0.02465753 | De Magalhaes and Costa 2009 |
| *Anser cygnoides* | NA | NA |
| *Meleagris gallopavo* | 0.03013699 | De Magalhaes and Costa 2009 |
| *Numida meleagris* | 0.02465753 | De Magalhaes and Costa 2009 |
| *Pavo cristatus* | 0.01369863 | De Magalhaes and Costa 2009 |
| *Phasianus colchicus* | 0.03013699 | De Magalhaes and Costa 2009 |
| *Parus major* | 0.0205 | Tomás et al. 2012 |
| *Ficedula albicollis* | 0.0178 | Gill and Donsker 2017 |
| *Corvus sp.* | 0.01068 | Holyoak 1967 |
| *Geospiza difficilis* | 0.0329 | Grant and Grant 1980 |
| *Taniopygia guttata* | 0.0151 | Olson et al. 2014 |
| *Rattus norvegicus* | 0.1003562 | De Magalhaes and Costa 2009 |
| *Microtus arvalis* | 0.0768 | De Magalhaes and Costa 2009 |
| *Microtus ochrogaster* | 0.04164384 | De Magalhaes and Costa 2009 |
| *Mus musculus musculus* | 0.104 | De Magalhaes and Costa 2009 |

| | | |
|---|---|---|
| *Mus spretus* | NA | NA |
| *Drosophila melanogaster* | 6.3 | Hanson et al. 1929 |
| *Drosophila teissieri* | NA | NA |
| *Drosophila santomea* | NA | NA |
| *Drosophila yakuba* | NA | NA |
| *Drosophila simulans* | NA | NA |
| *Drosophila sechellia* | NA | NA |

| **Species** | **Longevity (years)** | **source** |
|---|---|---|
| *Formica fusca* | 20 | Personnal communication |
| *Formica sanguinea* | 20 | Personnal communication |
| *Formica cunicularia* | 20 | Personnal communication |
| *Formica pratensis* | 6 | Personnal communication |
| *Melanargia galathea* | 1 | Lafranchis et al. 2015 |
| *Maniola jurtina* | 1 | Lafranchis et al. 2015 |
| *Aphantopus hyperantus* | 1 | Lafranchis et al. 2015 |
| *Pyronia tithonus* | 1 | Lafranchis et al. 2015 |
| *Pyronia bathseba* | NA | NA |
| *Mytilus californianus* | 25 | Bayne and Bayne 1976 |
| *Mytilus trossulus* | 25 | Bayne and Bayne 1976 |
| *Mytilus galloprovincialis* | 25 | Bayne and Bayne 1976 |
| *Mytilus edulis* | 25 | Bayne and Bayne 1976 |
| *Allolobophora chlorotica L1* | 1.25 | Edwards & Bohlen 1996 |
| *Allolobophora chlorotica L2* | 1.25 | Edwards & Bohlen 1996 |
| *Allolobophora chlorotica L4* | 1.25 | Edwards & Bohlen 1996 |
| *Aporrecta icterica* | NA | NA |
| *Lumbricus terrestris* | NA | NA |
| *Lineus lacteus* | NA | NA |
| *Lineus longissimus* | NA | NA |
| *Lineus sanguineaus* | NA | NA |
| *Lineus ruber* | NA | NA |
| *Homo sapiens* | 123 | De Magalhaes and Costa 2009 |
| *Pan troglodytes* | 59.4 | De Magalhaes and Costa 2009 |
| *Gorilla gorilla* | 60.1 | De Magalhaes and Costa 2009 |

| | | |
|---|---|---|
| *Papio anubis* | 37.5 | De Magalhaes and Costa 2009 |
| *Pongo abelii* | 59 | De Magalhaes and Costa 2009 |
| *Macaca mulatta* | 40 | De Magalhaes and Costa 2009 |
| *Anas platyrhynchos* | 29.1 | De Magalhaes and Costa 2009 |
| *Anser cygnoides* | 31 | De Magalhaes and Costa 2009 |
| *Meleagris gallopavo* | 13 | De Magalhaes and Costa 2009 |
| *Numida meleagris* | NA | NA |
| *Pavo cristatus* | 23.2 | De Magalhaes and Costa 2009 |
| *Phasianus colchicus* | 27 | De Magalhaes and Costa 2009 |
| *Parus major* | 15.4 | De Magalhaes and Costa 2009 |
| *Ficedula albicollis* | 9.8 | De Magalhaes and Costa 2009 |
| *Corvus sp.* | 19.2 | De Magalhaes and Costa 2009 |
| *Geospiza difficilis* | 9 | Oschadleus et al. 2016 |
| *Taniopygia guttata* | 12 | De Magalhaes and Costa 2009 |
| *Rattus norvegicus* | 3.8 | De Magalhaes and Costa 2009 |
| *Microtus arvalis* | 4.8 | De Magalhaes and Costa 2009 |
| *Microtus ochrogaster* | 5.3 | De Magalhaes and Costa 2009 |
| *Mus musculus musculus* | 4 | De Magalhaes and Costa 2009 |
| *Mus spretus* | NA | NA |
| *Drosophila melanogaster* | 0.16 | Linford et al. 2013 |
| *Drosophila teissieri* | NA | NA |
| *Drosophila santomea* | NA | NA |
| *Drosophila yakuba* | NA | NA |
| *Drosophila simulans* | NA | NA |
| *Drosophila sechellia* | NA | NA |

### Table S5: Values and sources of the life history traits used in this study.

*One of the two estimates of the adaptive rate at group level used in this study, $\omega_{a[A]}$, was obtained by calculating the across-species arithmetic mean of $\omega_{na}$ within a group, then subtracting this average from ω. Here we show that $\omega_{a[A]}$ is an unbiased estimate of the adaptive rate with fluctuating population size if the pace of fluctuations is sufficiently slow. We justify this strategy by making the hypothesis that the present variation in population sizes between closely related species represents well the possible range of population size fluctuations that one population experienced during the time period of its divergence with its sister species. Under our model, $\omega_{na}$ for an individual species is*

*(1)*

$$\omega_{na} = (\widehat{D_N^{na}}/L_N)/(D_S/L_S),$$

*where $\widehat{D_N^{na}}$ is the expected number of non-synonymous substitutions:*

*(2)*

$$\widehat{D_N^{na}} = 2L_N N_e t\mu \int_{-\infty}^{0} \phi(s) f(N_e, s) ds$$

*where $L_N$ is the number of non-synonymous sites, t is the divergence time, $\phi(s)$ is the fixation probability of a mutation with a selection coefficient s, and $f(Ne, s)$ is the DFE, in a population of size $N_e$.*

*Then, if we have sampled n closely related species, and we call $N_{e1}$, $N_{e2}$, $N_{e3}$, … $N_{en}$ their respective present effective population size, then, making the hypothesis that the shape of the DFE and the mutation rate μ remains constant in time, one can express the expected number of non-synonymous substitutions, $\widehat{D_N^{na}}$, as :*

*(3)*

$$\widehat{D_N^{na}} = \frac{t}{n} \sum_{i}^{n} 2L_N N_{ei}\mu \int_{-\infty}^{0} \phi(s) f(N_{ei,s}) ds$$

*Equation (3) is equivalent to assuming that, as the considered species were diverging, they have randomly fluctuated between the n regimes of selection/drift we currently observe, spending the same amount of time in the n regimes. Under this assumption, the group level $\widehat{D_N^{na}}$ is simply the arithmetic mean of $\widehat{D_N^{na}}$ estimated across individual species. Then, using $L_N$, $L_S$ and $D_S$ of the total subtree of the considered species, we can use the arithmetic mean of $\omega_{na}$ across individual species, $\omega_{na[A]}$, as representative of the average non-adaptive selective regime during their divergence. Subtracting $\omega_{na[A]}$ from the dN/dS ratio estimated using all branches of the tree, we obtain an estimate of the adaptive substitution rate for the whole group, $\omega_{a[A]}$.*

**Box S1: Rationale of the estimation of the per group adaptive substitution rate "A".**

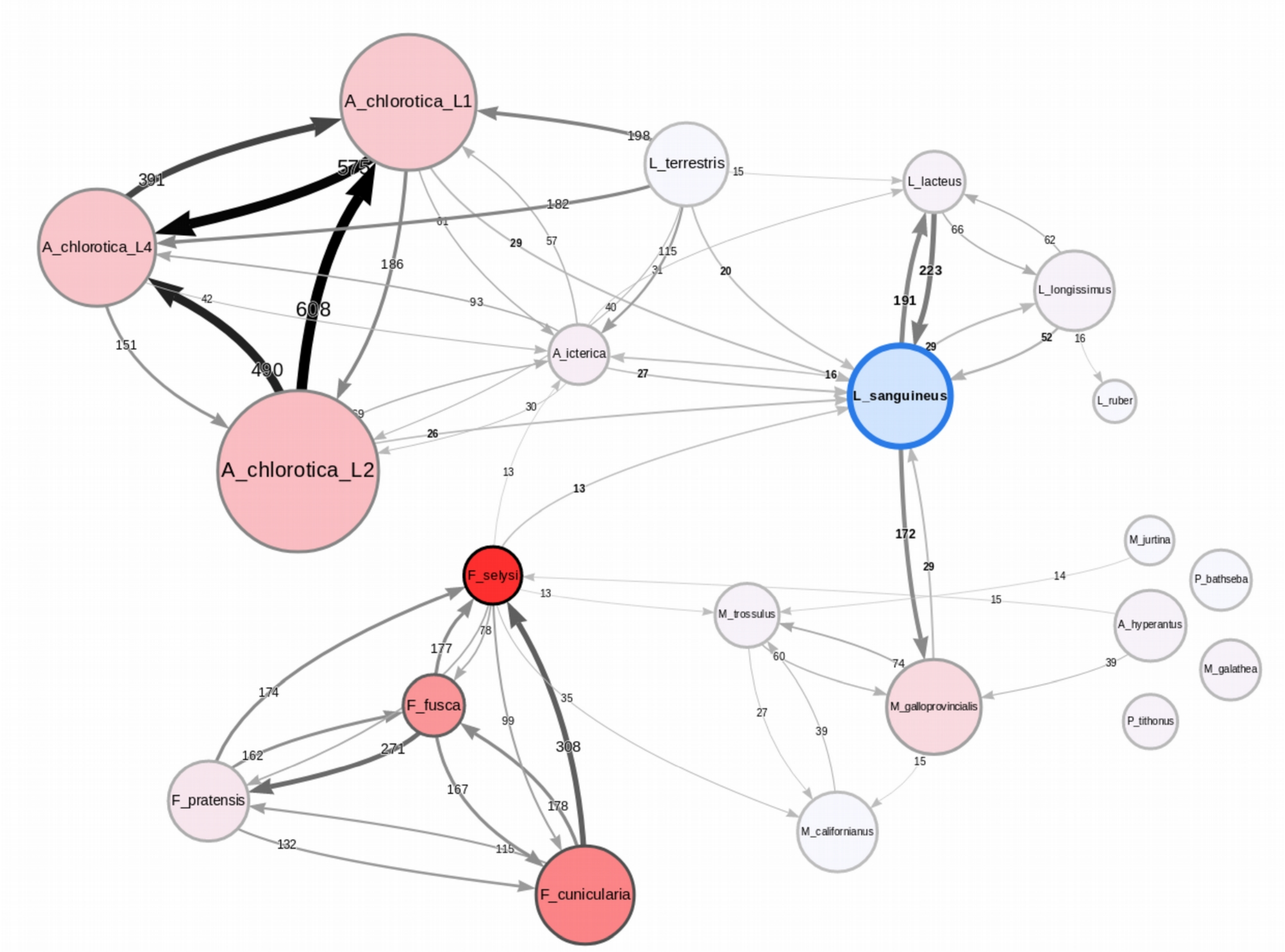


**Figure S1: Cross contamination network for de novo assemblies from exon capture.**

Circles represent the assemblies, and arrows and their corresponding numbers represent the number of cross contaminants. Most cross contamination events occur between closely-related species and are therefore likely false positive cases.

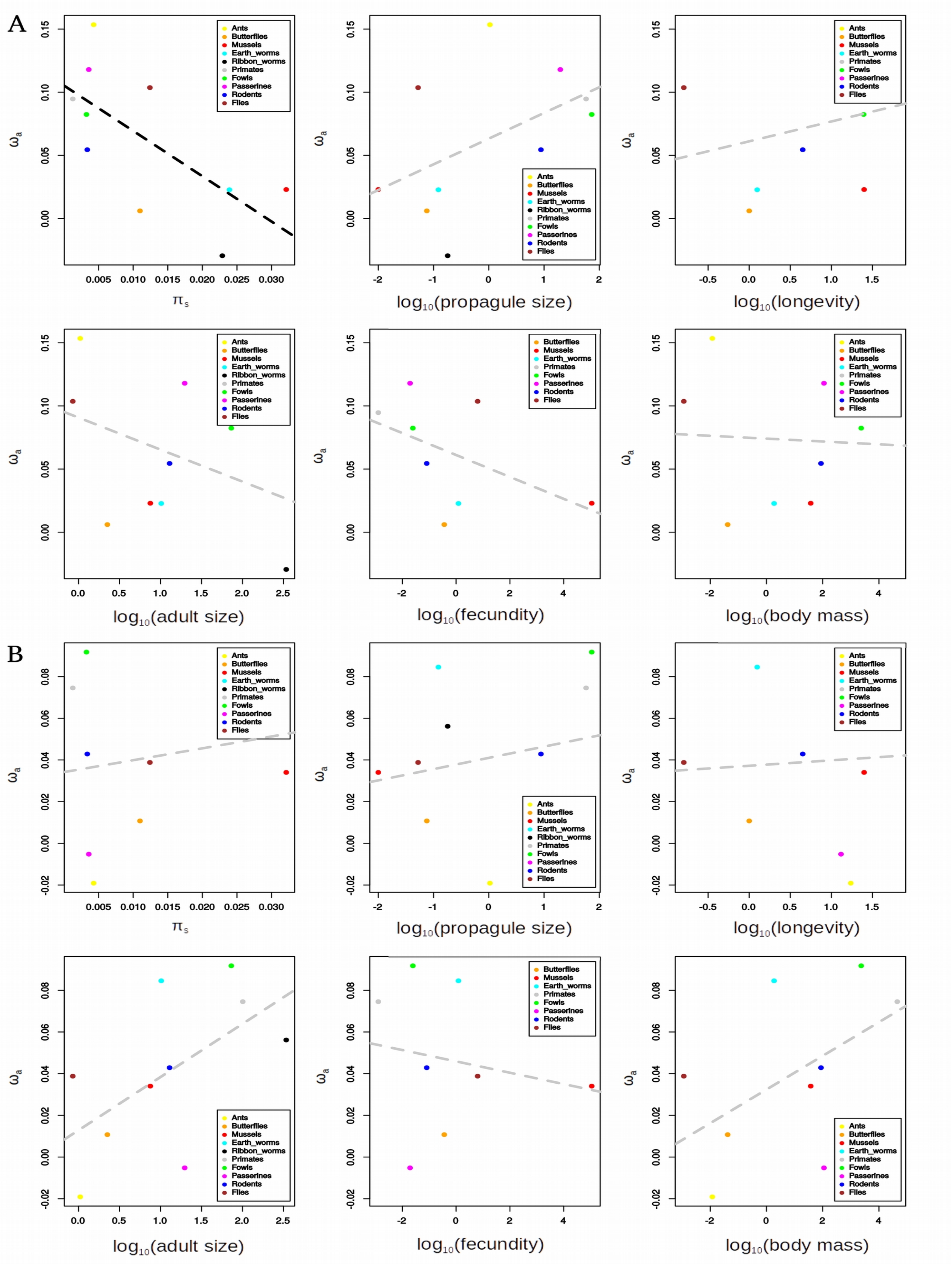


**Figure S2: Relationship between $\omega_{a[P]}$ and $\pi_s$ and $\log_{10}$ transformed life history traits.**

$\omega_{a[P]}$ is estimated using all mutations and substitutions (A) or using only GC-conservative mutations and substitutions (B). Group level $\pi_s$ and life history traits are estimated by averaging species level estimates across closely related species. Black dotted lines represent significant regressions across taxonomic groups and grey dotted lines non-significant ones.

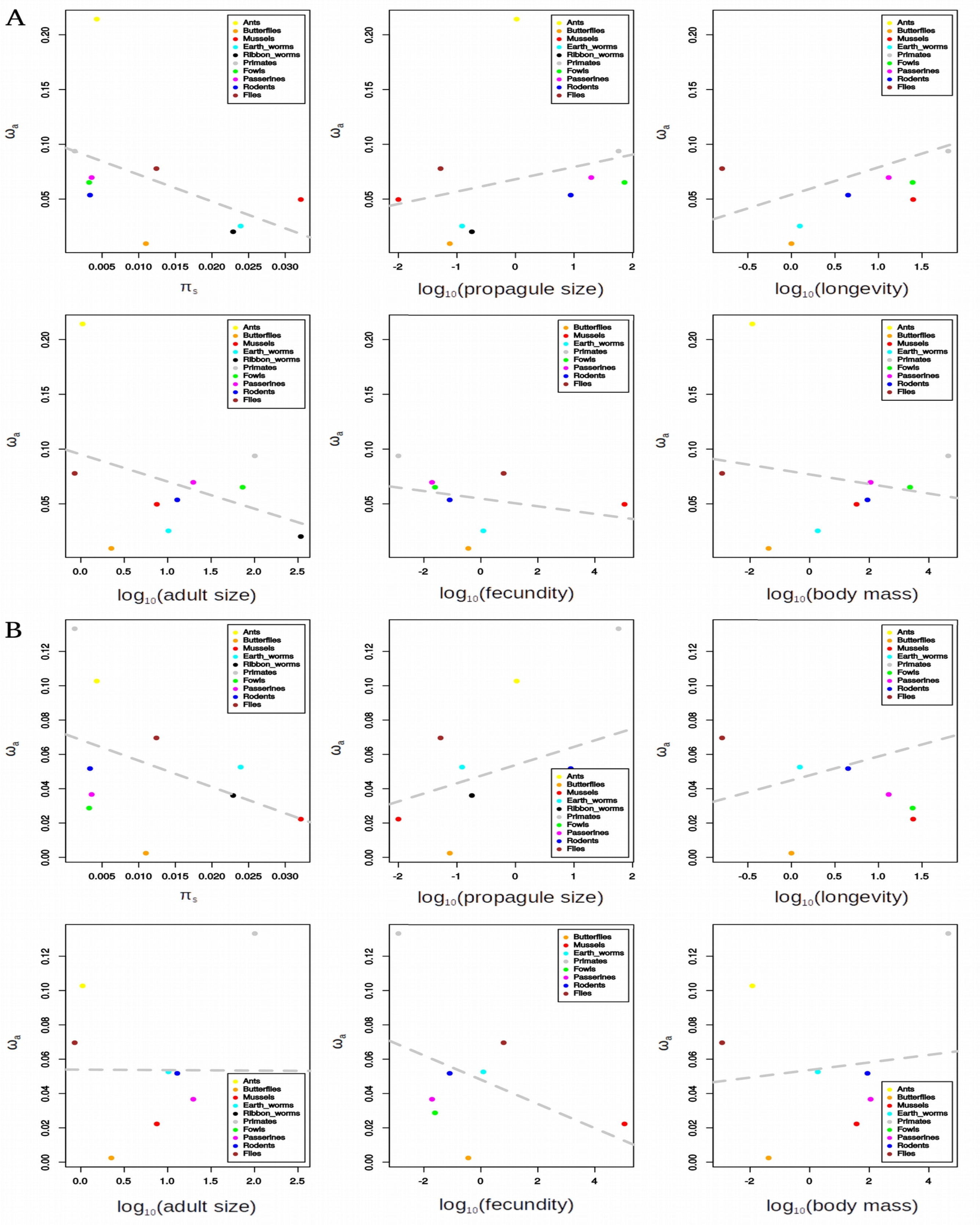


**Figure S3: Relationship between $\omega_{a[A]}$ and $\pi_s$ and $\log_{10}$ transformed life history traits.**

$\omega_{a[A]}$ is estimated using all mutations and substitutions (A) or using only GC-conservative mutations and substitutions (B). Group level $\pi_s$ and life history traits are estimated by averaging species level estimates across closely related species. Black dotted lines represent significant regressions across taxonomic groups and grey dotted lines non-significant ones.

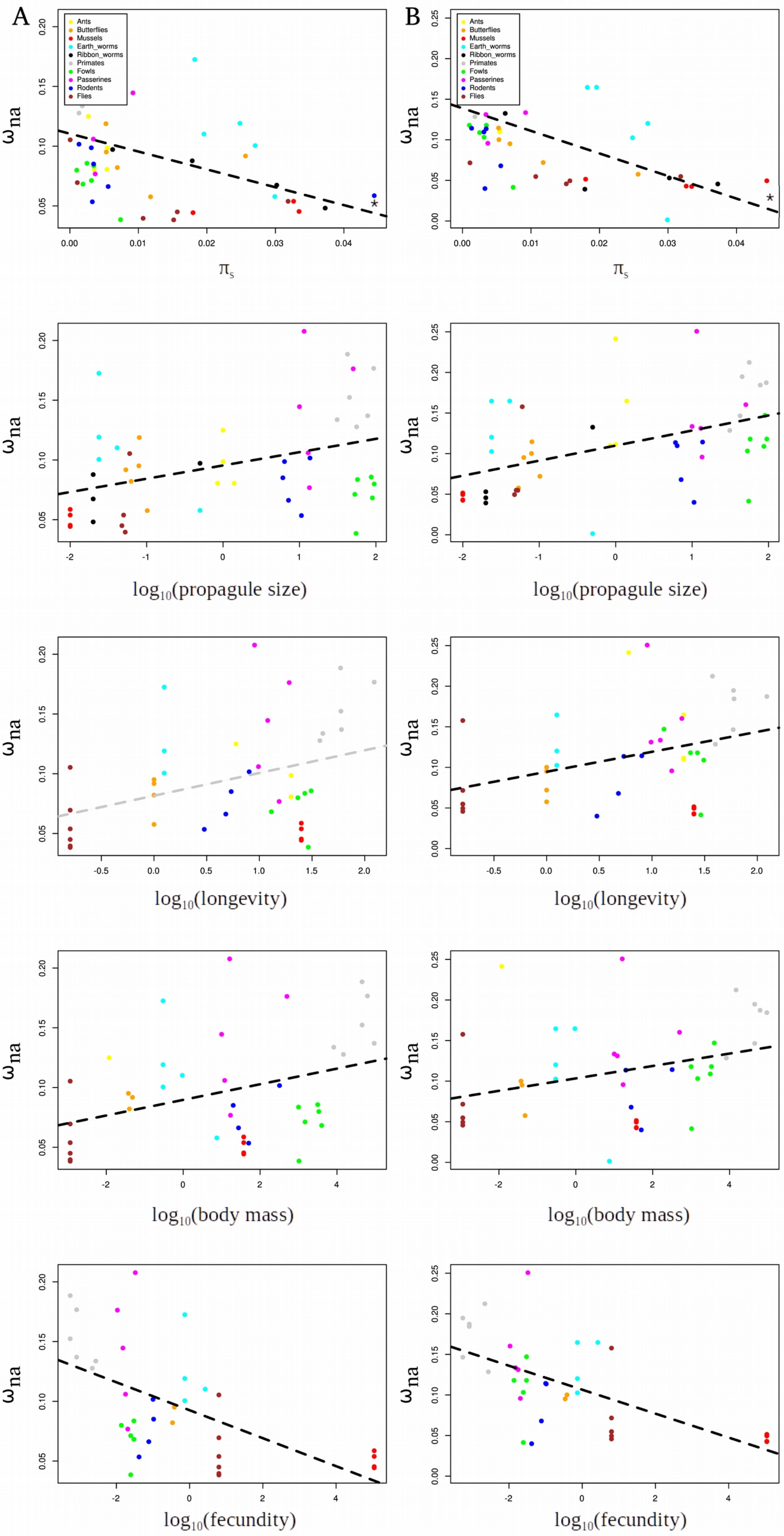


**Figure S4: Relationship between species-level $\omega_{na}$ and $\pi_s$ or $\log_{10}$ transformed life history traits.** $\omega_{na}$ is estimated using all mutations and substitutions (A) or using only GC-conservative mutations and substitutions (B). Black dotted lines represent significant regressions across taxonomic groups and grey dotted lines non-significant ones.

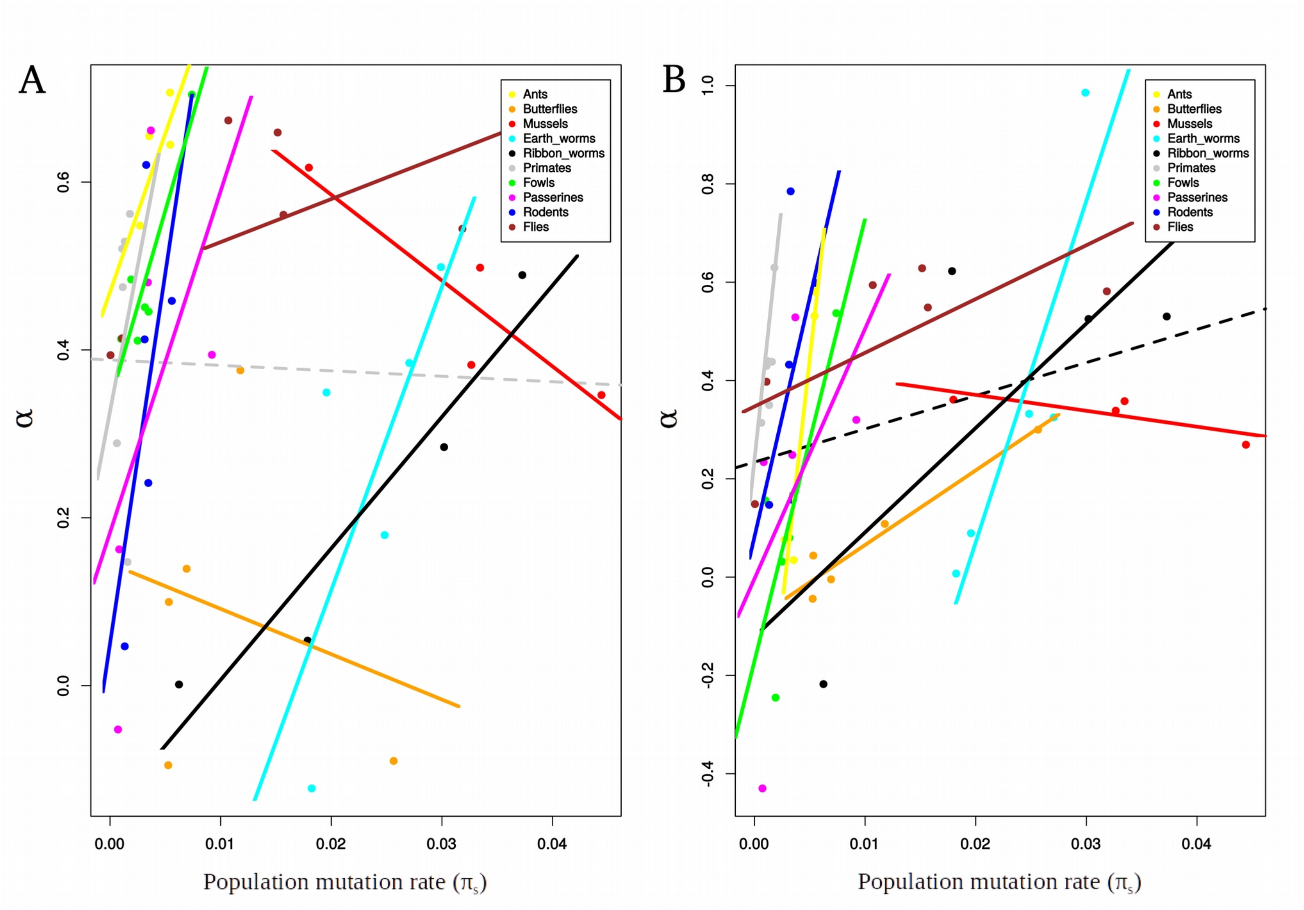


**Figure S5: Relationship between species-level α and $\pi_s$.**

*α* is estimated using all mutations and substitutions (A) or using only GC-conservative mutations and substitutions (B). The dotted line represents the regression across all species, and full lines represent the regression within each taxonomic groups. Black dotted lines represent significant regressions across taxonomic groups and grey dotted lines non-significant ones.

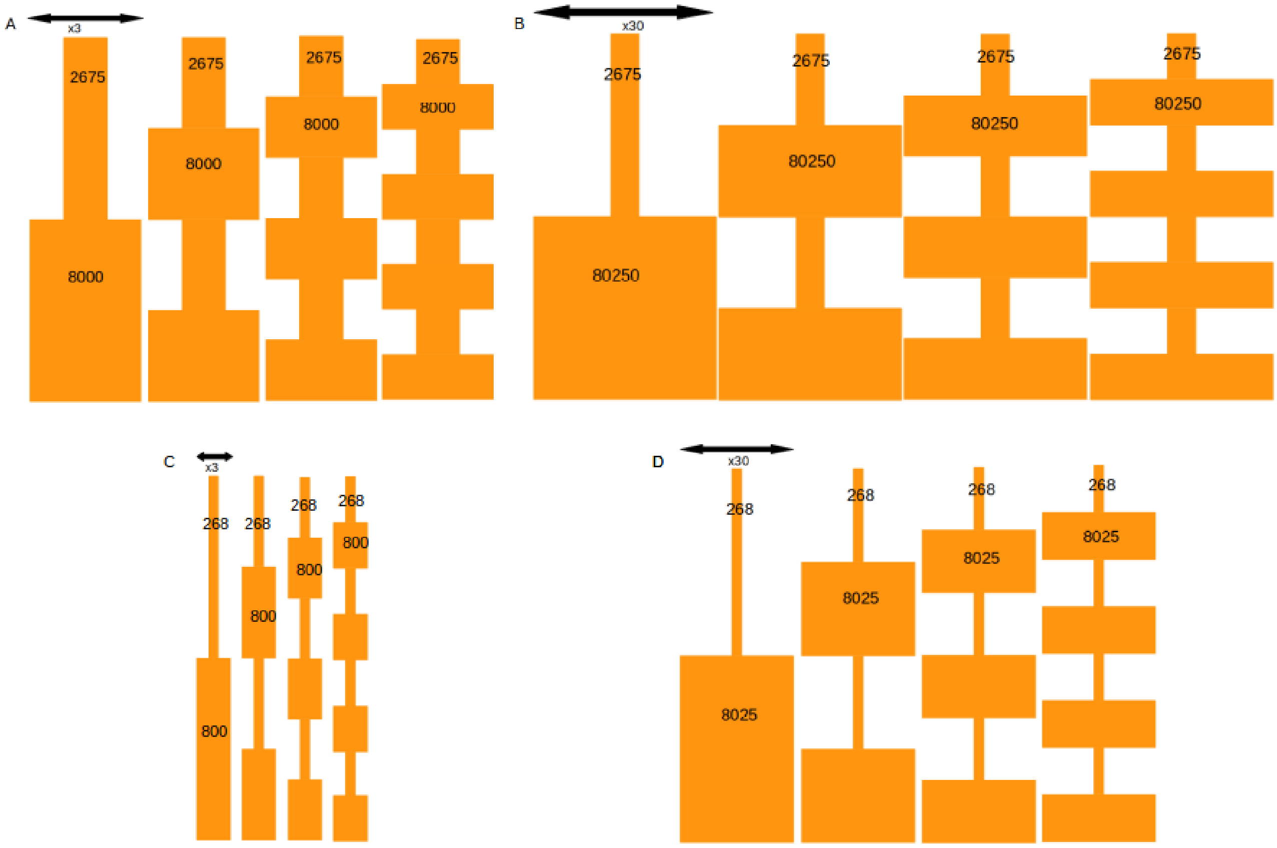


**Figure S6: Design of the simulations of fluctuation of population size.**

A: three fold ratio between low and high population size and high long-term population size.

B: thirty fold ratio between low and high population size and high long-term population size.

C: three fold ratio between low and high population size and low long-term population size.

D: thirty fold ratio between low and high population size and low long-term population size.

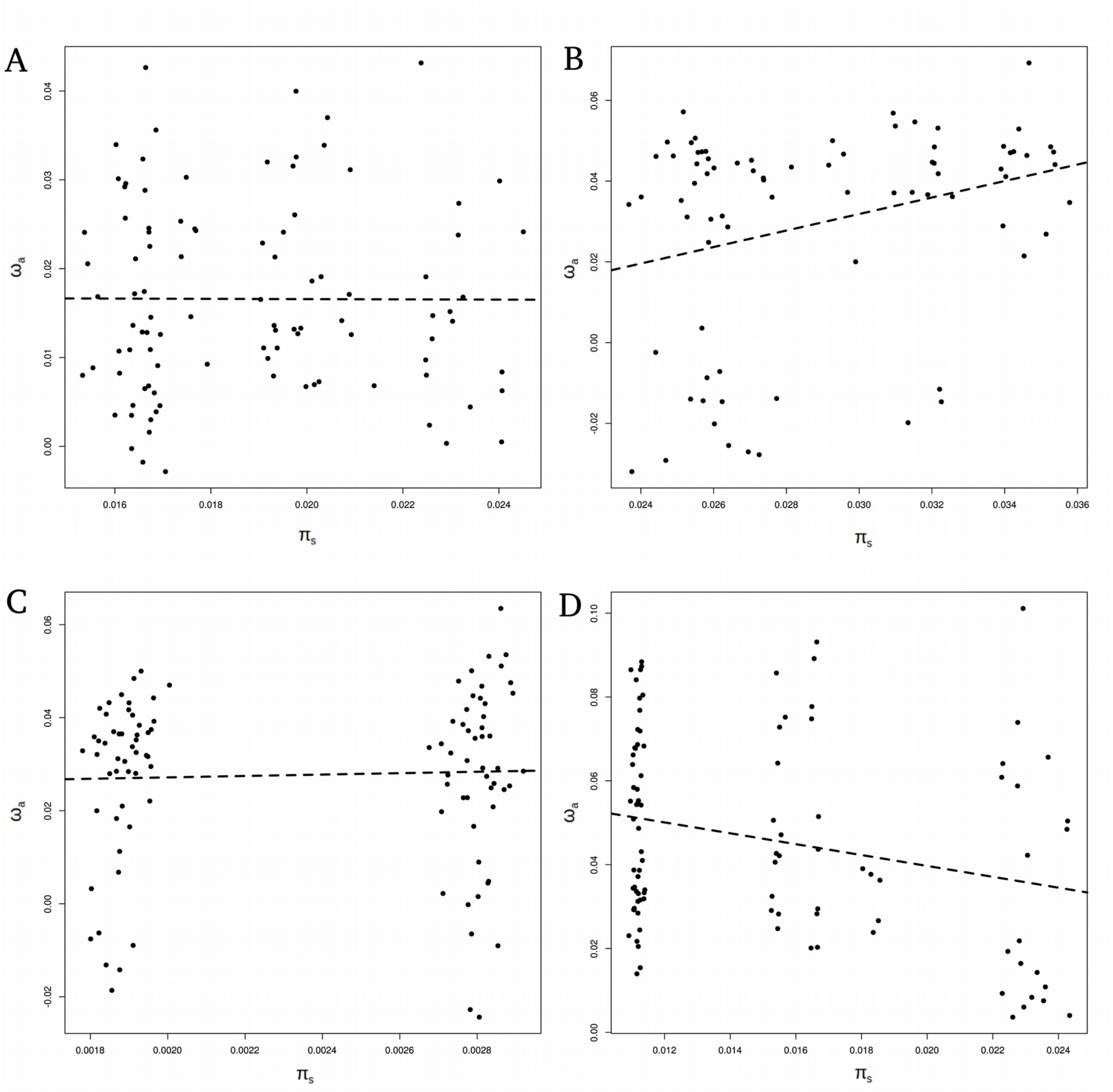


**Figure S7: Relationship between $\omega_a$ and $\pi_s$ in simulated scenarios of fluctuating population size.**

A: three fold ratio between low and high population size and high long-term population size (scenario A in figure S1)

B: thirty fold ratio between low and high population size and high long-term population size (scenario B in figure S1)

C: three fold ratio between low and high population size and low long-term population size (scenario C in figure S1)

D: thirty fold ratio between low and high population size and low long-term population size (scenario D in figure S1)
